# Life history traits mediate elevational adaptation in a perennial alpine plant

**DOI:** 10.1101/2023.10.17.562199

**Authors:** Aksel Pålsson, Ursina Walther, Simone Fior, Alex Widmer

## Abstract

- Spatially divergent natural selection drives adaptation to contrasting environments and the evolution of ecotypes. Understanding this process in perennial plants is challenging because natural selection acts on multiple life history traits linked by fitness trade-offs.
- In a multi-year reciprocal transplant experiment of high and low elevation populations of the alpine carnation *Dianthus carthusianorum* in the Central Alps, we tested how different stages of the life cycle contribute to adaptation. Moreover, we used matrix population models to infer the specific contributions of individual life stages to fitness, coupled with trade-off analyses.
- We found genotype x environment interactions consistent with elevational adaptation both in single fitness components linked to reproduction and survival, and in integrative fitness estimates. Adaptation at low elevation is driven by early reproduction, in contrast to an opposite strategy at high elevation. Adaptive life-history differences between populations originating from low and high elevations are mediated by environmental effects on plant growth and trade-offs between reproduction and survival.
- Our work reveals elevational ecotypes of the perennial alpine plant *D. carthusianorum* that express alternative life history strategies in response to climatic differences shaping resource allocation.

## Introduction

Spatially divergent natural selection drives adaptation of populations to contrasting environmental conditions (Savolainen *et al*., 2013) and may lead to the evolution of phenotypically divergent ecotypes (Turesson, 1922; Wadgymar *et al*., 2022). Adaptation in plants is well studied owing to their amenability to reciprocal transplant experiments, where fitness, the contribution of an individual or genotype to the next generation, can be quantified in different environments. In annual species, fitness is commonly estimated through seed output, as this captures life-time reproductive success as determined by selection acting on traits expressed in one growing season (Ågren & Schemske, 2012; Bischoff & Hurault, 2013). In perennial species, on the other hand, reproduction is conditional to selection acting in subsequent stages of a multi-year life cycle, where each stage provides a complementary component of life-time fitness (Kim & Donohue, 2011a; DeMarche *et al*., 2020; Wadgymar *et al*., 2017). This complicates the measure of fitness, as the inherent costs of each life-stage may imply trade-offs with others. As a result, populations growing in different environments may evolve life history traits that optimize the allocation of resources at different stages of the life cycle to maximize life-time fitness (Stearns, 1992; Friedman, 2020; Forbis & Doak, 2004, Shefferson *et al*., 2003). Thus, assessing the impact of selection across the life cycle and dissecting how the interactions between fitness components shape the evolution of life history traits is key to understanding adaptation in perennial plants.

Life history traits of perennial plants, such as age- or size-specific reproduction and longevity, characterize the investment of individuals in separate fitness components (Stearns, 1992; Laiolo & Obeso, 2017; Acasuso-Rivero *et al*., 2019). Life history theory predicts interactions between the expression of individual fitness components that depend on resource allocation (Stearns, 1992; Friedman, 2020; Hamann *et al.,* 2021). In resource-rich environments, maximization of one fitness component may have positive effects on others. In most natural environments, however, resources are limited and their allocation is typically associated with trade-offs. A classic example is investment into reproduction, which may lead to reduced future survival and growth (Obeso, 2002; Sletvold & Ågren, 2015; Hamann *et al*., 2021). In general, trade-offs can vary across the life span of perennial plants and can be exacerbated under adverse environmental conditions (Stearns, 1992; Acerenza, 2016; Hamann *et al*., 2021; von Euler *et al*., 2012). Hence, contrasting selection pressures in different environments can lead to the evolution of ecotypes expressing divergent strategies (Stearns, 1992; Childs *et al*., 2010: Friedman *et al*., 2020; Boyko *et al*., 2023). Characterizing the adaptive role of life history traits requires long-term field experiments where fitness is modelled in an integrative framework to assess stage-specific contributions. Yet, despite the importance of life history traits in the evolution of perennial plants, this is rarely done.

Matrix population models (MPMs) offer a powerful analytical framework to estimate population growth rates by integrating fitness and trade-offs across multiple fitness components of the life cycle (Caswell, 2001). When applied to multi-year reciprocal transplant experiments, MPMs provide an integrative estimate of population fitness suitable to compare performance of alternative populations growing in the same environment. Moreover, life-table response experiments (LTRE) are analytical methods applied to MPMs which can identify vital rates, i.e., stage or age-specific fitness components, with the strongest contributions to adaptation (Caswell, 1989). The influence of specific vital rates on population growth are expressed as elasticities. Age classes that compose each population are further estimated as stable age distributions (Caswell, 2001). Together, these estimates can provide complementary descriptors of the impact of different life history traits on the responses of populations to different environments. Even though MPMs are a powerful tool for life history analysis, they remain underutilized in adaptation studies (Wadgymar *et al*., 2022; but see e.g., Waser & Price, 1985; DeMarche *et al*., 2016; Goebl *et al.,* 2022), because they require comprehensive datasets obtained from multi-year experiments and ideally include seedling establishment, a stage of critical importance for plant fitness but relatively rarely assessed in the field (Kitajima & Fenner, 2000; Kim & Donohue, 2011a).

Understanding the contributions of life history traits to fitness is particularly important for species that are predicted to face substantial environmental change, such as alpine plants (Anderson & Song, 2020; Tito *et al*., 2020; Lancaster *et al*., 2016). Plant populations growing along elevational gradients typically experience contrasting environmental conditions, and often show evidence for local adaptation (Halbritter *et al*., 2018). Ongoing warming is exacerbated in mountain ranges, and altered selection through both abiotic and biotic factors may threaten future population persistence (Nomoto & Alexander, 2021; Alexander *et al*., 2015; Bemmels & Anderson, 2019). Plant life history traits often vary along elevational gradients, which suggests that different strategies may be advantageous under contrasting environmental conditions (Laiolo & Obeso, 2017). While this variation may have a genetic basis, it may also result from phenotypic plasticity, the ability of genotypes to express different phenotypes in different environments (Price *et al*., 2003; Ghalambor *et al*., 2007; Acasuso-Rivero *et al*., 2019). Considerable plasticity has been documented in life history traits across a wide range of organisms (Davidson *et al*., 2011; Palacio-Lopez *et al*., 2015; Ensing & Eckert, 2019), and while there is no consensus on the extent to which plasticity influences adaptation, plasticity can provide an immediate response to changing conditions to support transient population persistence (Fox *et al*., 2019; Nicotra *et al*., 2010; Matesanz *et al*., 2010).

Here, we test whether populations of the perennial alpine carnation *Dianthus carthusianorum* are adapted to the contrasting environmental conditions at low and high elevation in the Alps, and assess the contributions of life history traits to this process through a multi-year reciprocal transplant experiment. Specifically, we tested genotype by environment (GxE) interactions driven by selection using individual fitness components at subsequent stages of the life cycle, and combined this evidence with an experimental test of seedling establishment to generate MPMs. Using this experimental framework, we asked: 1) Has selection driven the evolution of elevational ecotypes, and which stages of the life cycle impact adaptation? 2) How do life history traits affect plant performance in contrasting environments, do they imply fitness trade-offs, and how do they contribute to the adaptation process? 3) How do contrasting environmental conditions alter the expression of specific vital rates of populations of alternate elevational origin? We hypothesize that climate-driven selection linked to elevation has driven adaptation mediated by life history traits as a result of trade-offs involved in plant performance. We further hypothesize that environmental variation impacts the contribution to population growth of vital rates expressed by high and low elevation populations.

## Materials and methods

### Study system

*Dianthus carthusianorum* L. (Caryophyllaceae) grows on dry and nutrient poor grasslands and rocky slopes at elevations from sea level up to the alpine zone above 2’100 m asl in the Alps (Landolt, 1985; GBIF 2022). It is a short-lived herbaceous perennial with a woody taproot and a basal rosette of grass-like leaves. Upon flowering, it produces one to several mostly unramified generative stalks, each with a single terminal inflorescence consisting of up to fifteen pink to purple flowers (Garcke, 1972). *D. carthusianorum* is gynodioecious, self-compatible but primarily outcrossing, and main pollinators are diurnal butterflies (Bloch *et al*., 2006).

Our study was performed in the in the Upper Rhône Valley (Valais, Switzerland) in the central Alps (Figure 1), a major east-west oriented inner-Alpine valley with a dense yearly precipitation distribution leading to a particularly dry climate at low elevation (Braun-Blanquet, 1961; Zumbrunnen *et al*., 2009). Erosion during Pleistocene glaciations generated a high relief landscape with peaks over 4000 m (Sternai *et al*., 2013), resulting in pronounced climatic gradients between dry and warm conditions at low elevation, and wetter and cooler wet conditions at higher elevation. In September 2015, we established two common gardens each at low and high elevation (~900 and 2100 m, respectively; Table S1). We established a reciprocal transplant experiment with populations originating from opposite ends of the elevational distribution range of *D. carthusianorum* in the Upper Rhône Valley. Three low (i.e.,~700-900 m) and three high (i.e., ca. ~2000-2200 m) elevation populations were sampled in neighboring valleys to represent the species’ distribution in the colline and lower alpine belts (Table S2). Each transplant site was equipped with a weather station (DS3 IP66, SensorScope, Lausanne, Switzerland). Comparison of climate parameters with long term long term records (i.e., 1980-2018) from the Chelsa database (Karger *et al*., 2017; Karger *et al*., 2021) indicate that the transplant sites capture the different climate experienced by the six populations at their original sites (Figure S1). Warmer absolute temperatures measured at the present time relative to the time series average may be attributed to a 0.05(°C) annual increase over the past 38 years (Figure S2). Soil potential measured at a depth of 30 cm further highlights the summer drought that affects the low elevation sites (Figure S3).

**Figure 1.**
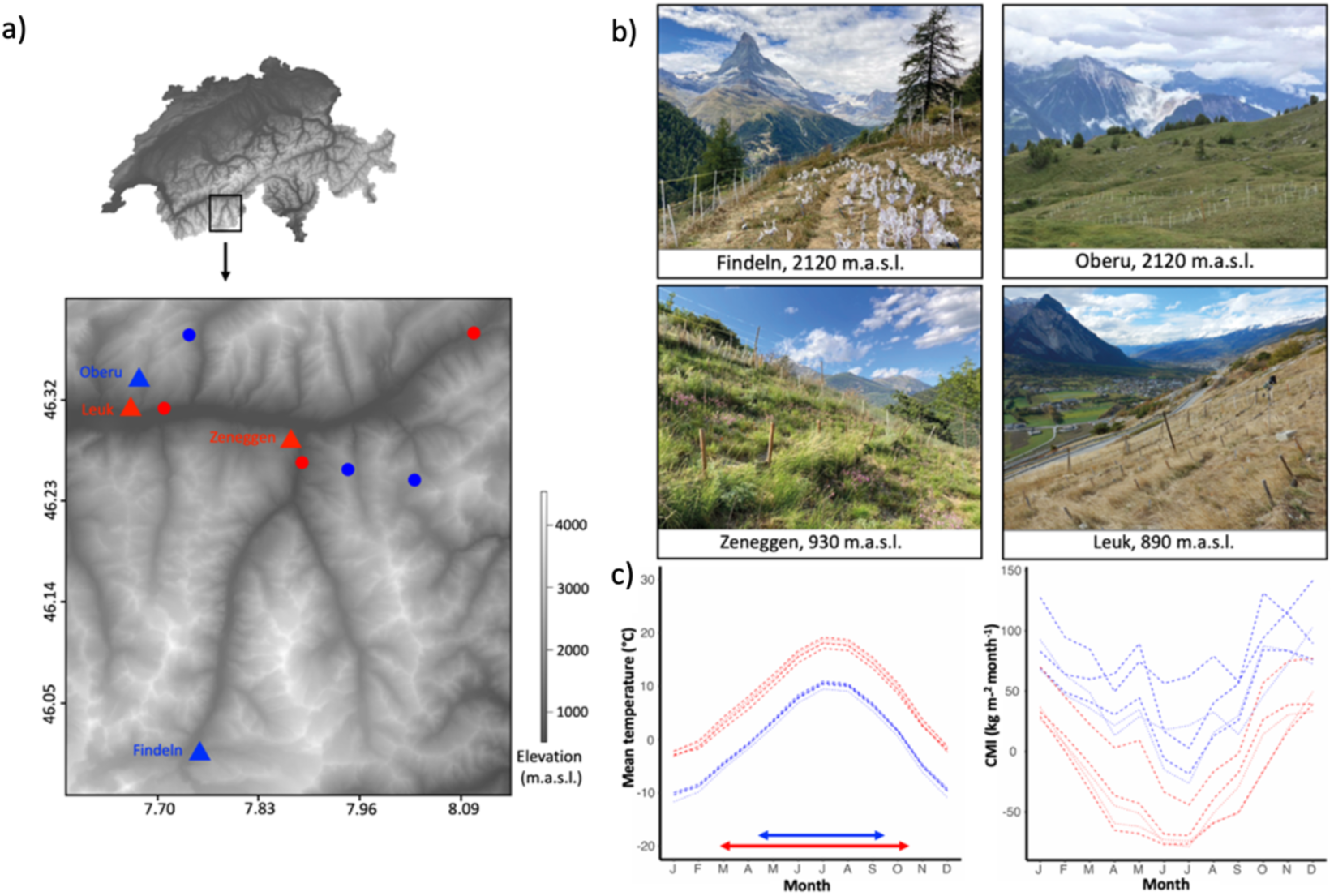
Experimental set up of reciprocal transplant experiments. a) Locations of the sampled *D. carthusianorum* populations (circles) and the transplant sites (triangles) in the study area in the Upper Rhône Valley (Switzerland). b) Low (bottom panels) and high elevation (top panels) transplant sites, c) Mean monthly temperature and Climate moisture index, CMI at low (red) and high (blue) elevation transplant sites (dotted lines) and at the original sites of the wild populations (dashed lines). CMI indicates the difference between amount of precipitation and potential evapotranspiration, negative values are hence associated with dry conditions. Estimates are the monthly averages calculated over the timespan 1980-2018 using the Chelsa high-resolution data base (Karger *et al.,* 2021). The arrows indicate the growing season.

### Reciprocal transplant experiment

We collected seeds from 20-39 plants (i.e., families) in each of the six focal populations in fall 2012 and 2014. In summer 2015, we germinated the seeds and grew seedlings over a period of three months in a greenhouse at ETH research station Lindau-Eschikon (Switzerland) in peat moss based soil (Klasmann Deilmann Gmbh) under a 12-hour day/night cycle, with temperatures set to 20 and 18°C during the day and night, respectively, and a relative humidity of 50-60%. In fall 2015, we transplanted ~500 seedlings from 127-135 families at each of four transplant sites (Table S1, Table S2). Seedlings were placed in eight randomized blocks, approximately 1 m apart. Each block contained 72 individuals, arranged in a 12×6 grid and spaced 25 cm apart. As the start of winter following the onset of the experiment was exceptionally warm and dry, we watered seedlings twice a week until mid-January to help establishment. Individuals that died within two weeks after being transplanted were attributed to transplant shock and excluded from the analyses. The sites were fenced to exclude large herbivores commonly present in the study area, such as deer, cattle, and marmots. Throughout the experiment, we trimmed the surrounding vegetation to prevent the experimental plants from becoming overgrown.

### Assessment of vital rates

We monitored vital rates (i.e., age specific survival and flowering probability, and seed count) over the growing seasons 2016-2018, encompassing three reproductive transitions. At the low sites, we defined the start and end of each growing season as the first and last date of 6-day windows with mean daily temperatures above 10 °C. Because the weather stations had to be removed during winter at the high sites to prevent damage from snow, we defined the start of the growing season based on the observed start of vegetative growth, which approximately corresponded to three weeks after snowmelt. The growing seasons ranged from March to November at low elevation, with flowering occurring typically in June, and from late May to October at high elevation, with flowering primarily in July (Figure 1). We recorded survival at the start and end of each growing season and visited each site twice a week in 2016 and once a week in 2017 and 2018 to record flowering individuals. We bagged individual inflorescences in organza bags for seed collection after flowers had wilted. In the laboratory, we extracted seeds from capsules by carefully separating undeveloped seeds and quantified seed output as the average number of mature seeds from two independent runs of an elmor C3 High Sensitive Seed Counter (Elmor Ltd, Schwyz, Switzerland). We estimated the size of the basal rosette at the start and end of each growing season (except the end of the second year) from high resolution pictures of each plant (Nikon D810; 7360×4912 pixels) including a reference standard, and by computing the mean of orthogonal diameters measured in image J v.2.0 (Schindelin *et al.,* 2012). This estimate of plant size provides a suitable proxy for biomass as evidenced by its correlation with plant dry weight (Figure S4).

### Seedling establishment

We assessed germination and seedling establishment success of all six focal populations growing in each transplant site for implementation in MPMs. From the seed harvested in 2017 at each site, we selected ~200 seeds from each populations represented by fruiting plants belonging to up to five families originally collected in the wild populations. At the high elevation sites, only seeds from one low and two high elevation populations could be included in the experiment due to low seed production by individuals from the other populations. In fall 2018, we sowed ~10 seeds per family in the same sites in which they were produced, using peat moss soil (Klasmann Deilmann Gmbh) contained in biodegradable pots of 10×8 cm. We placed 96-165 pots per site in the ground after random assignment to positions within the blocks of the original experiment left empty by dead individuals. Plants that germinated successfully and were alive at the end of the 2019 growing period were considered established.

## Statistical analyses

### Separate vital rates

We assessed elevational adaptation from separate vital rates including flowering probability, seed output and survival, and tested for differential plant size. We tested genotype by environment interactions, as well as for differences between genotypes growing within each environment (local vs foreign criterion for local adaptation) and for the effect of the environment on each genotype (home vs away criterion, Kawecki & Ebert, 2004). We fitted separate generalized linear mixed effect models for flowering probability, seed output and survival at subsequent stages of the life cycle. In each model, we implemented the vital rate as response variable and genotype, transplant environment and their interaction as predictors, using a binomial error distribution for the categorical variables (i.e., flowering probability and survival), and a zero-inflated Poisson error distribution for the count variable (i.e., seed output). We analysed plant size using the same model structure as above, but implemented in linear mixed effect models with a Gaussian error distribution and we log transformed the data to improve distribution of the residuals.

In all analyses, we implemented families nested within population and block nested within site as random effects. We implemented mixed effect and zero-inflated models using the R package lme4 v1.1 (Bates *et al*., 2015) and glmmTMB v. 1.3 (Brooks *et al*., 2017), respectively, and used DHARMa v. 0.4.6 to perform model diagnostics of the zero-inflated models (Hartig, 2022). We assessed significance levels of interactions using likelihood ratio tests. We obtained estimates, significance levels and confidence intervals of the contrasts between genotypes with the emmeans package (Lenth, 2017). We analysed survival throughout the life cycle using mixed effect cox models, which perform proportional hazards regression of time to event data with implementation of random effects. We fitted the Cox models with genotype and transplant environment and their interaction as predictors and used the same random effect structure as used in the mixed effect models in the package survival 2.44 (Therneau & Grambsch, 2000). All analyses were performed in R v.3.3.2 (R Core Development Team 2016).

### Integrated fitness estimate

We formulated age-structured MPMs to obtain an integrated estimate of fitness expressed as population growth rate *λ* for each genotype growing at low and high elevation. To obtain survival and reproductive vital rates, we divided the plant life-cycle into winter (W_i_) and summer (S_i_) stages (Figure S5). We calculated the survival vital rates as the proportion of individuals transitioning (T_i_) to the next life stage. As an integrated value of the reproductive vital rates (R_i_) we used the product of flowering probability, seed count and recruitment. From the seedling experiment, we estimated the establishment rates per genotype in each site from generalized linear mixed effect models by implementing the proportion of seedlings established per pot as the response variable and genotype as the predictors, with binomial error distribution and the same random effects structure as described above for the single vital rate analyses. We recovered similar establishment rates of the two genotypes conditional on the growing environment, with overall higher values in the low (i.e., low genotype: 0.17 ± 0.091; high genotype: 0.14 ± 0.073; Table S3) compared to the high elevation environment (i.e., low genotype: 0.02 ± 0.028; high genotype: 0.06 ± 0.058; Table S3). To assess whether differences in population growth rates between the genotypes growing in each environment are statistically significant, we performed 20 000 bootstrap replicates of each matrix stratified by population, and constructed bias corrected 95% confidence intervals around estimates. To decompose the effects of specific vital rates on population growth rate, we used life-table response experiments (LTRE) (Caswell, 1989). LTREs tests the difference in *λ* between a matrix of interest and a reference matrix, breaking down the contributions to adaptation from individual vital rates. We compared the matrix of the foreign genotype against the matrix of the local genotype in each environment. All MPM analyses were performed using packages popbio v. 2.2.4 (Stubben & Milligan, 2007) and boot v. 1.3 (Canty & Ripley, 2021).

### Fitness trade-offs

To examine how trade-offs contribute to adaptation, we modelled specific vital rates as the response variable of a three-way interaction between the following predictors: a vital rate at a preceding stage of the life cycle, genotype, and growing environment. Specifically, we tested for trade-offs between the probability of flowering and plant size at the start of the same growing season; the probability of survival and plant size at the start of the same growing season; and the probability of winter survival and flowering probability in the previous growing season. We used generalized linear models with binomial error distribution to test for three-way interactions, as well as two-way interactions between vital rates and the genotype within the two environments. We determined the significance of the interactions using likelihood ratio tests and computed pairwise contrasts between genotypes as well as confidence intervals of the trends using the R package emmeans (Lenth, 2017).

### Elasticity analyses and stable age distributions

To elucidate the influence of individual life history traits on population growth of the genotypes growing in the low and high elevation environments, we extracted the elasticities of specific vital rates and stable age distributions from the MPMs. The elasticities estimate the proportional sensitivities of a change in a specific vital rate to population growth rate, thus yielding their relative importance. As an outcome of trade-offs between survival and reproduction, stable age distributions describe the proportion of the population that is found in each age class at equilibrium. Hence, the elasticities of genotypes growing in the alternative environment reflect estimates of the environmental effect on the importance of specific vital rates for population growth rates (Caswell, 2001). We compared estimates at different stages of the life cycle by computing 95% bias corrected confidence intervals from 20 0000 bootstrap replicates of the matrix models. To further describe the environmental effect on the elasticity values, we correlated the shift (i.e, difference in value when growing in a home vs. a foreign environment) and distance (i.e., difference in value between the two genotypes growing in their home environments) of the three populations representing each genotype. We refrained from assessing significance of the correlations because elasticities are non-independent parameters derived from the population growth models.

## Results

### Evidence for adaptation from separate vital rates

We found significant GxE interactions for flowering probability in each year (Figure 2a-c, Table S4). The local genotype had significantly higher flowering probability at low elevation the second and third year, and at high elevation in the third year, thus fulfilling the local vs. foreign criterion. The low genotype also performed significantly better at home than away across the three years.

**Figure 2.**
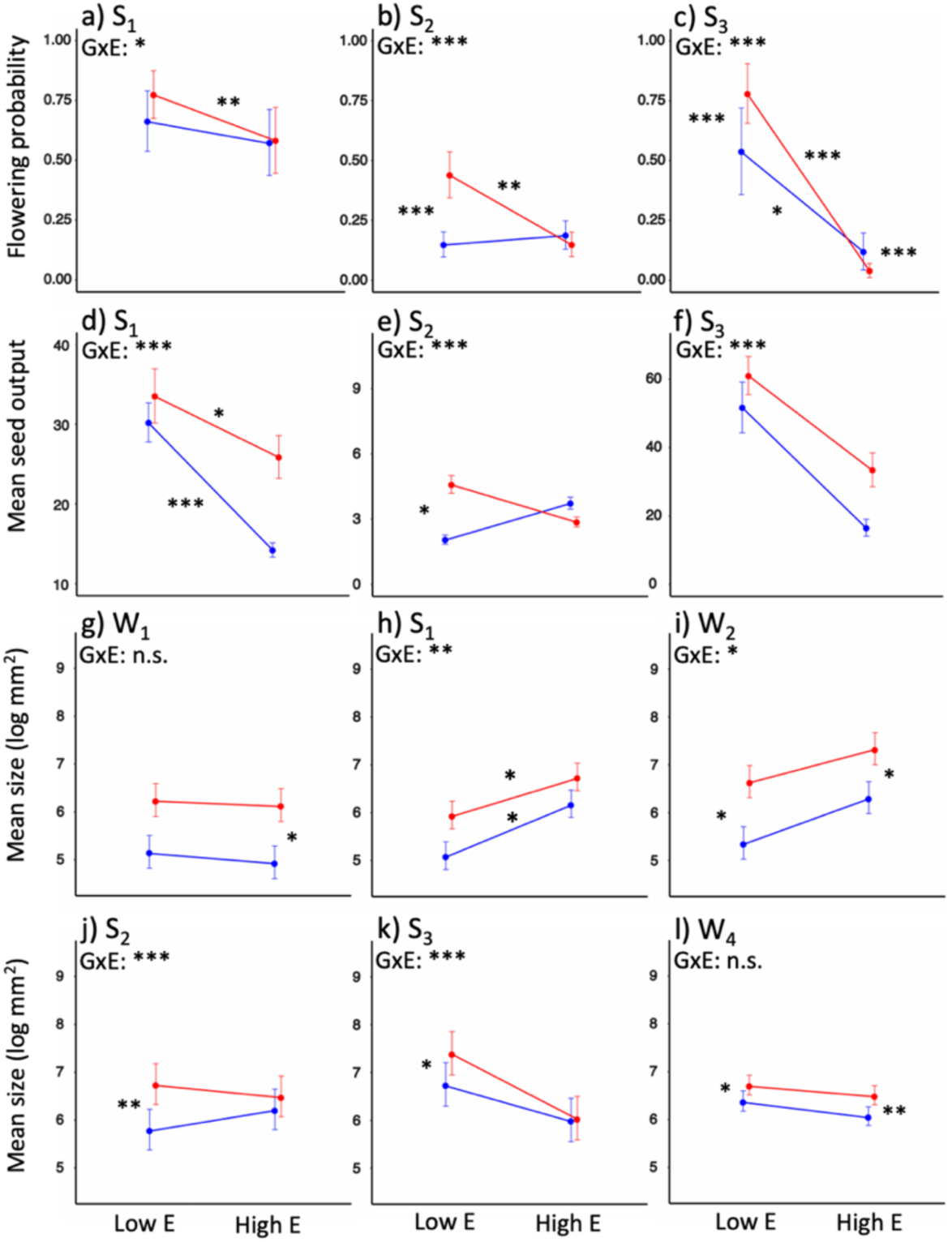
Reaction norms of genotypes for reproduction (a-f) and plant size (g-l) at subsequent stages of the life cycle. Symbols indicate mean estimate values inferred from mixed effect models and bars indicate 95% confidence intervals. Mean values are connected by reaction norms depicting the effect of the environment on each genotype. Red and blue colors denote the low and high genotypes, respectively. S_i_ and W_i_ denote summers and winters, respectively, of subsequent years. Significance of GxE interactions and contrasts consistent to the local vs. foreign and home vs. away criteria are reported (***p<0.001, **p<0.01, *p<0.05).

Reaction norms for seed output recapitulated patterns observed for flowering probability. Significant GxE interactions in each year (Figure 2d-f, Table S4) were associated with a fitness advantage of the local genotype at low elevation the second year. This genotype also showed a home vs. away advantage the first year and non significant trends in a consistent direction the following years.

Reaction norms for plant size varied substantially across the different stages of the plant life cycle (Figure 2g-l, Table S5). Significant GxE interactions were found at the start of each vegetation period (S_1_-S_3_), and over the second winter (W_2_). During the entire duration of the experiment, the low genotype was consistently larger than the high elevation counterpart, even when growing at high elevation, though differences were not always statistically significant.

Cumulative survival was significantly higher in the local genotype in both environments (Figure 3, Table S6) with a stronger cumulative effect at low elevation. While the low genotype showed similar cumulative survival at both elevations, the high genotype performed significantly better in its home environment (Table S6). The most pronounced differences of the survival curves between genotypes were observed at alternative time points in the two environments. At low elevation, relative survival rates of the local genotype were significantly higher than the foreign genotype during the first two summer seasons (S_1_ and S_2_), with a particularly pronounced difference during the first (Figure 3, Table S7). In contrast, at high elevation, survival rates of the local genotype were significantly higher than the foreign genotype during the first and second winter (W_1_ and W_2_; Figure 3, Table S7).

**Figure 3.**
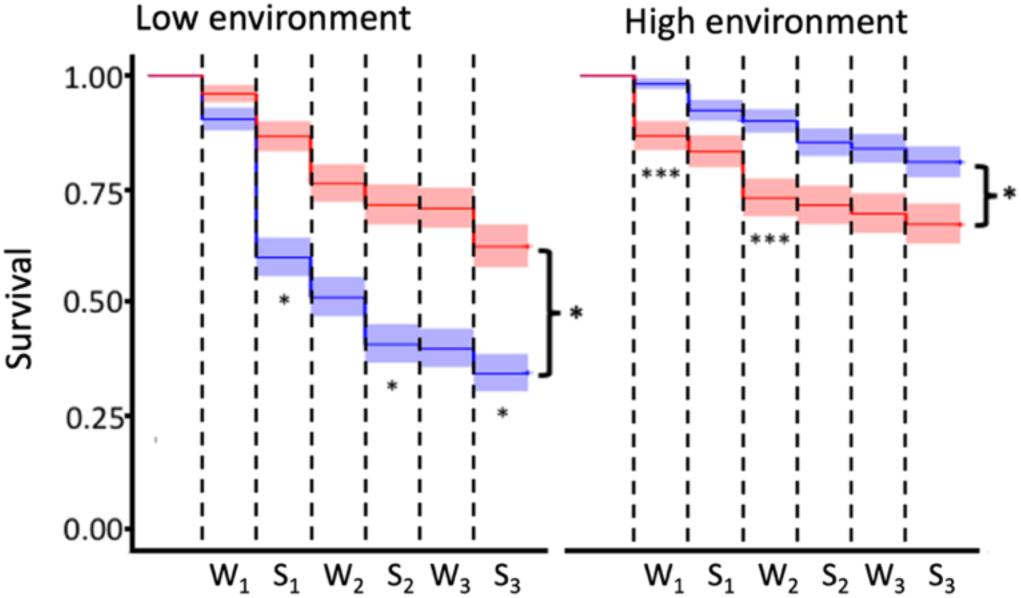
Survival of the genotypes throughout the experiment at subsequent stages of the life cycle. Red and blue colors denote the low and high genotypes, respectively. Survival curves of the two genotypes are inferred from cox proportional hazard models, with shaded areas representing 95% confidence intervals. Asterisks at the side of each plot indicate significant cumulative differences between the genotypes. Asterisks underneath the curves indicate significant differences of survival probability between genotypes at specific life stages, as assessed by generalized linear mixed effect models (***p<0.001, **p<0.01, *p<0.05).

### Evidence for adaptation from the integrated fitness estimates

Comparisons of population growth rates inferred from MPMs revealed a significant advantage of the local genotype in both the low and high elevation environment, with *λ* of the local genotype surpassing that of the foreign genotype by 35% and 57%, respectively (Figure 4, Table S8). LTRE analyses showed a negative impact of the foreign genotype on vital rates underlying population growth, predominantly driven by reproduction over survival (Figure 4, Table S8). Notably, differences in population growth were largely driven by reproduction in the first year in the low environment. At high elevation, the impact of individual vital rates was more evenly spread, with the strongest difference observed in the third year (R_3_).

**Figure 4.**
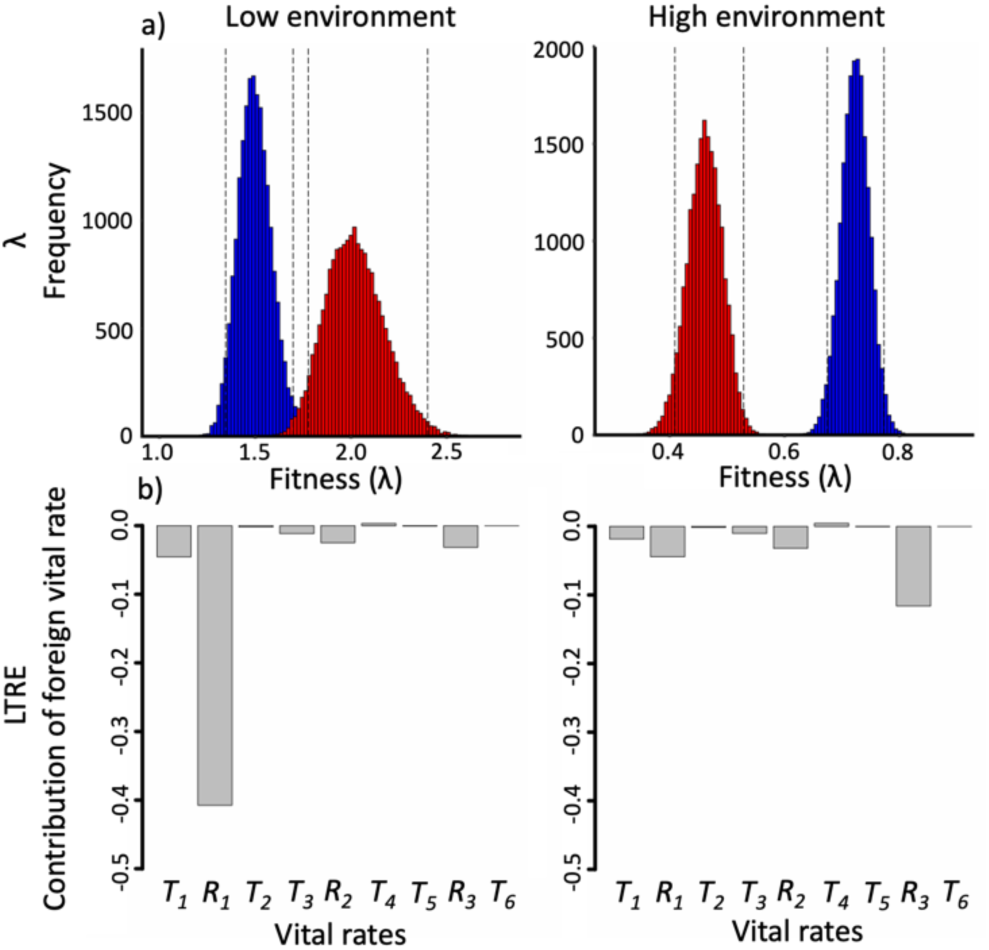
Integrative estimates of fitness. a) Histograms representing population growth rate distributions based on 20 000 bootstrap replicates. Dotted lines indicate bias corrected 95% confidence intervals. Red and blue colors denote the low and high genotypes, respectively. b) LTRE showing the relative contribution to population growth of vital rates expressed as survival and reproduction (T_i_ and R_i_, respectively, with _i_ indicating the age specific vital rates) of the foreign genotype at subsequent stages of the life cycle.

### Fitness trade-offs

*i) Size and flowering probability.* The flowering probability varied as function of a significant three-way interaction among plant size at the start of summer, genotype and environment in the first and second year (Figure 5a, d; Table S9). Flowering probability consistently increased with increasing plant size, though with a stronger effect in the high compared to the low genotype in both environments.
*ii) Size and survival*. Plant size generally showed a positive effect on survival throughout the experiment, with significant effects at multiple stages of the life cycle (Table S10). These effects were particularly relevant during the first year of flowering and subsequent winter. In the former season, we recovered a significant three-way interaction, and larger size was generally associated with higher survival (Figure 5b, c, Table S10). However, the high genotype showed contrasting effects on survival in the first summer (S_1_) and the following winter (W_2_), as larger plants suffered lower survival in the latter season (Figure 5e, f, Table S10).
*iii) Flowering probability and survival*. Survival varied as a function of a significant three-way interaction among flowering probability, genotype and environment at the end of the first summer and the following winter. In the former season, flowering had a significant positive effect on survival of each genotype in each environment (Table S11). Conversely, in the latter it negatively affected the survival of the high genotype in the low elevation environment (Figure 5g; Table S11). Moreover, low elevation plants that flowered in the high elevation environment suffered higher mortality relative to the local flowering plants (Figure 5g: Table S11).

**Figure 5.**
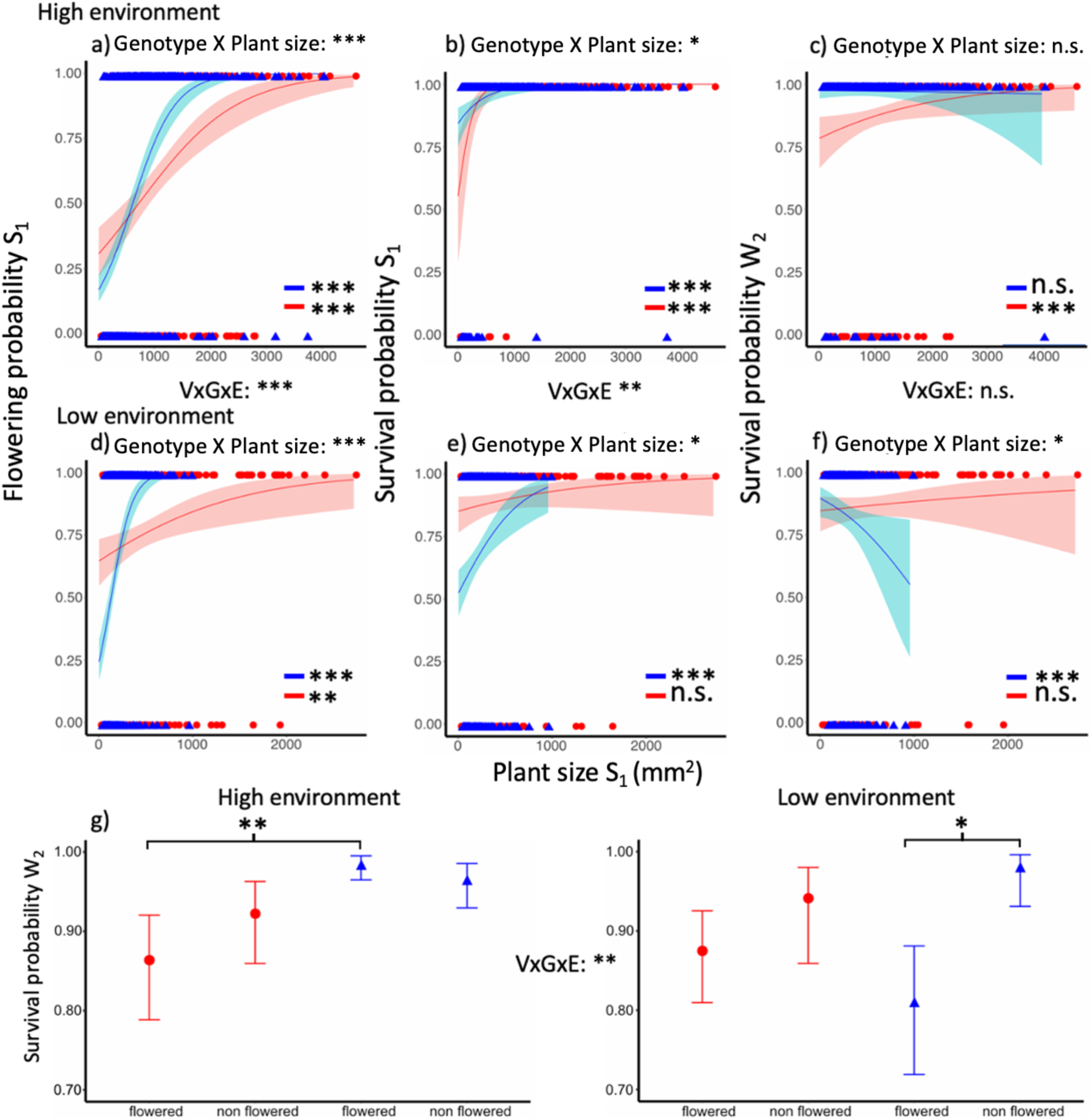
Fitness trade-offs expressed by the genotypes. a-f) The effect of size on flowering probability the first summer (S_1_), survival probability the first summer and second winter (W_2_) Lines indicate predicted relationships from generalized linear model regressions with 95% confidence intervals. Symbols indicate empirical values of plant size and flowering probability. Red and blue colors denote the low and high genotypes, respectively. Significant relationships are indicated within each panel as asterisks. g) The effect of flowering the first year on survival probability the second winter (W_2_). Symbols indicate mean estimate values inferred from generalized linear models and bars indicate 95% confidence intervals. Red and blue colors denote the low and high genotypes, respectively. In all plots, for each vital rate used as response variable, significance of the three-way interaction between the vital rate used as predictor, the genotype and the transplant environment (VxGxE), as well as the two-way interaction between the vital rate used as predictor and the genotype within the low and high elevation environment is reported (***p<0.001, **p<0.005, *p<0.05).

### Elasticities and stable age distributions

Elasticity values showed divergent patterns describing the influence of subsequent vital rates on population growth (Figure 6a, b, Table S12). Survival during the first winter (T_1_) and reproduction the first summer (R_1_) had by far the strongest influence at low elevation. In comparison, vital rates were more evenly spread at high elevation. Notably, elasticity values of both genotypes were similar within environments, with non significant differences at each stage of the life cycle. Similarly, stable age distributions differed substantially between environments, but showed similar trends between genotypes within environments. At low elevation, populations consisted primarily of young individuals, with a marked shift to adults at high elevation (Figure 6c, d, Table S13). Additionally, we observed correlations between elasticity shifts (i.e., difference in elasticity of age specific vital rates when growing a home vs. foreign environment) and distance (i.e., difference in elasticity of age specific vital rate between the two genotypes growing at the home elevation) for both genotypes (Figure 6e, f) in response to the high and low elevation environments.

**Figure 6.**
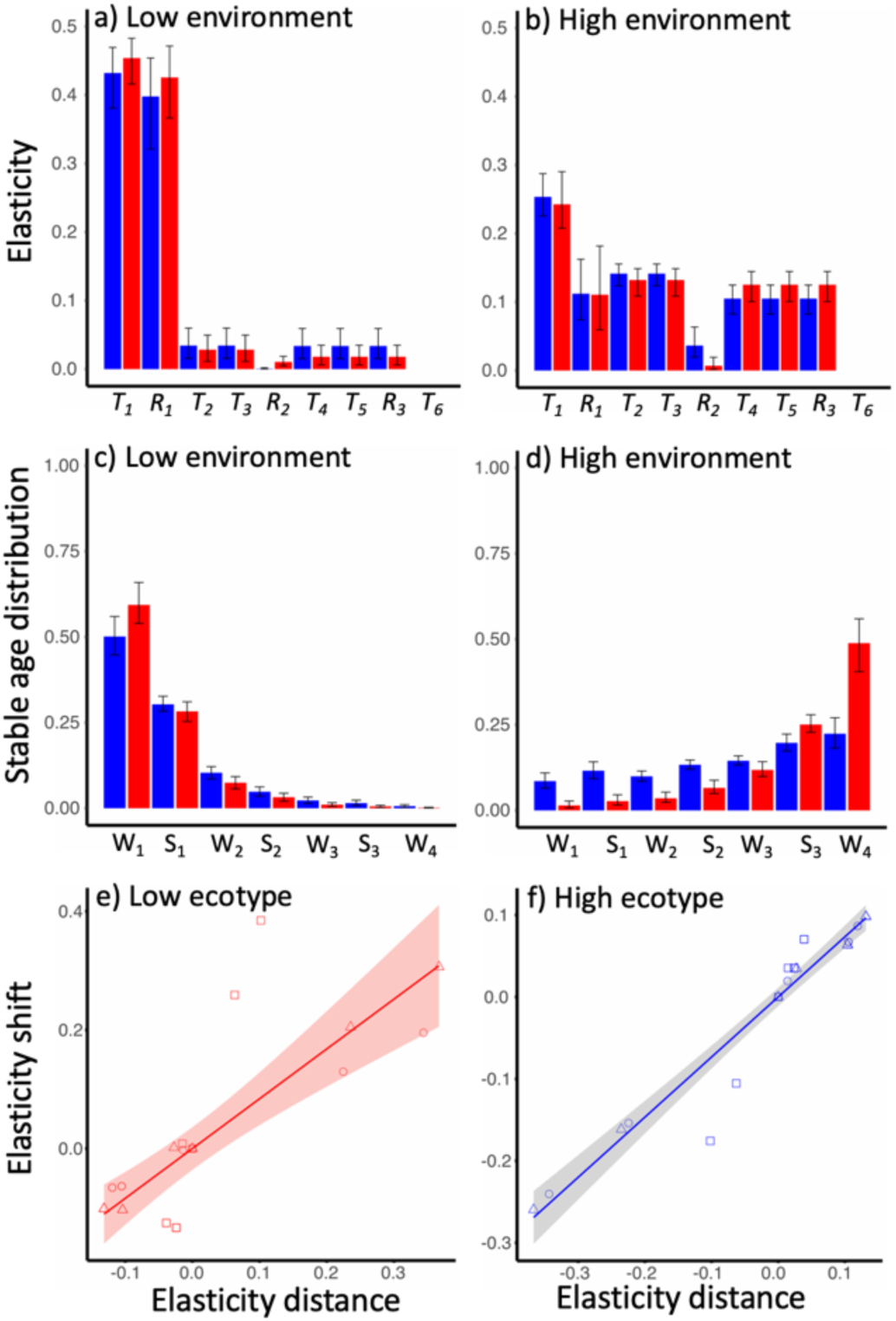
Elasticities and stable age distributions of the genotypes. a, b) Influence of the specific vital rates of survival and reproduction (T_i_ and R_i_, respectively, with _i_ indicating the age specific vital rates) on population growth rates. c, d) Stable age distribution at subsequent summer and winter (S_i_ and W_i_, respectively, with _I_ representing subsequent years) seasons. In a-d, red and blue bars indicate estimates for the low and high genotypes, respectively. e, f) Relationship between elasticity shift (i.e., difference in elasticity of age specific vital rates when growing at home vs. foreign environment) and elasticity distance (i.e., difference in elasticity of age specific vital rate between the two genotypes growing at the home elevation) expressed by the three focal populations (symbols) representative of each genotype growing in their respective home and away environment. The shaded area indicates 95% confidence intervals of the trendline.

## Discussion

Our multi-year reciprocal transplant experiment reveals that populations of *D. carthusianorum* originating from the lower and upper limits of the elevational distribution range in the central Alps are adapted to the contrasting environmental conditions and thus form elevational ecotypes (Turesson, 1922). Elevational adaptation is reflected by differences in single age-specific fitness components, as well as the population growth rate (*λ*), an integrated fitness estimate. Our recording of fitness across multiple life stages allowed us to infer the contributions of divergent life history traits to elevational adaptation. We found that high investment in fecundity of young individuals was advantageous at low elevation. This is a strategy that markedly differed from a more conservative investment in fecundity to the benefit of self-maintenance at high elevation. Moreover, the relative influence of life history traits to fitness was environment dependent, aligning with strategies expressed by locally adapted ecotypes. Our study emphasizes the importance of considering multi-year variation in performance linked to growth and development, life history traits and contrasting environmental conditions to understand adaptation of perennial plant systems.

Reproduction is often implicated as a key determinant of plant fitness (Hereford, 2009; Younginger *et al*., 2017). However, along elevational gradients, survival has been found to shape adaptation in many species (Halbritter *et al*., 2018). In this study, we found strong evidence for the contribution of both fitness components to elevational adaptation in *D. carthusianorum*. Flowering probability and seed output, our two fitness components related to reproduction, showed similar results. The signals of adaptation were stronger for the low genotype, particularly as it showed evidence of adaptation in flowering probability during the later years and consistent trends in seed output. These two fitness components are linked to bolting and resource allocation during seed set and are not functionally correlated (Angert & Schemske, 2007; Hautier *et al*., 2009), and thus contribute independently to adaptation. Survival was consistently higher for each genotype in its home environment. At low elevation, selection imposed on the foreign genotype was most pronounced during summers when plants were exposed to high temperatures and seasonal drought periods (Figure S3) that characterize the climate of our study region at colline elevational belt (Braun-Blanquet, 1961; Zumbrunnen *et al*., 2009). This corresponds to other systems where drought has been shown to be a key stressor (Kim *et al*., 2013; Orsenigo *et al*., 2014). In contrast, the high elevation environment imposed strong selection on low elevation plants during winter, most likely because of the depletion of resources during the extended snow cover. Hence, our results reveal that spatially divergent natural selection mediated by climatic differences affects both survival and reproductive fitness components and acts as driver of elevation adaptation.

While individual fitness components provide clear evidence of adaptation, further insights can be gained from our estimates of population growth rates (*λ*), as they account for multi-year contributions of individual vital rates of genotypes exposed to environmental selection. The absolute *λ* values should be interpreted with caution, in particular because populations were tested under conditions that are primarily representative of climatic differences between elevations, but only partially account for biotic components of selection (Cremieux *et al*., 2008; Hargreaves *et al*., 2020); however, relative comparisons provided compelling evidence of differential performance between genotypes. Results from LTREs further reveal that the contributions to adaptation in the genotypes are driven by early and late reproductive vital rates at low and high elevation, respectively. This evidence implicitly reflects the significant GxE interactions found in single fitness components, which individually constitute the vital rates implemented in MPMs. For both genotypes, vital rates expressed in a foreign environment are accompanied by trade-offs that are particularly evident between reproduction and survival. For example, flowering of the high genotype at low elevation expressed a cost reflected in reduced ability to survive during the following winter. Similarly, the low genotype flowering at high elevation suffers higher mortality relative to the local plants. Moreover, we infer that plant growth has a strong impact on both flowering and survival, and thus plays a key role as a determinant of trade-offs. Low elevation plants were consistently larger than high elevation plants regardless of the growing environment, and similar patterns in reaction norms across elevation indicate a consistent environmental effect on this trait across genotypes. Larger plants are overall more likely to flower but this effect is conditional to both the genotype and the environment, pointing to a genetic contribution to the interactions expressed across elevation. Similarly, larger size is overall beneficial for survival the first winter, but leads to a negative impact on high elevation plants growing in the low elevation environment in the following season. We interpret this evidence as a trade-off triggered by the environmental conditions in the low elevation environment which causes high elevation plants to exceeded allocation of resources to reproduction, thus compromising their physiological ability to survive the following winter. In line with other species occurring along elevational gradients, size in *D. carthusianorum* is thus better interpreted as a divergent phenotypic trait exhibiting spatially plastic variation that feedbacks on key fitness components rather than a fitness proxy *per se* (Bonser & Aarsen, 2009; Cheplick, 2020; Boyko *et al*., 2023; Halbritter *et al*., 2018; Körner 2003, Midolo *et al.,* 2020). Overall, our results suggest that adaptive genetic variation regulating trade-offs between fitness components of genotypes mediate adaptation in *D. carthusianorum*. Moreover, it appears that differential growth rates between genotypes that feedback on both reproduction and survival plays a primary role in this trade-off.

The allocation of resources to reproduction and survival is a fundamental mechanism governs adaptive life history strategies (Stearns, 1992). Our evidence of age specific fitness advantage of the local ecotypes in both reproductive and survival vital rates, point to divergent life history traits between the genotypes. Specifically, selection at low elevation favors the evolution of a life history strategy characterized by high investment in early reproduction, whereas at high elevation, selection favors self-maintenance over high reproduction. These inferences are congruent with expectations for plant systems inhabiting high energy environments, such as our low elevation habitats, characterized by abundant resources and intense competition, where early reproduction and high seed output are advantageous and compensate for high juvenile mortality (von Arx *et al*., 2006; Kim & Donohue, 2011a; Laiolo & Obeso, 2017; DeMarche *et al*., 2020;). Conversely, at high elevation, a life cycle characterized by reduced annual reproduction and allocation of resources to self-maintenance constitutes a better strategy under a short growing season and limited resources (Childs *et al*., 2010; Johnstons & Pickering, 2004; Milla *et al*., 2009). In fact, the alternative life history strategies expressed by our two genotypes at contrasting elevations are in line with observed differences in species occurring along elevation gradients, where short lived species with high reproductive investment are typical at lower elevations and long lived species with enhanced offspring quality and bet hedging strategies predominate at high elevation (Boyko *et al*., 2023; Laiolo & Obeso, 2017).

Elasticities are prospective estimates indicating the influence of each vital rate on population growth of each genotype, i.e., independent from a comparison with the other (Caswell 2000). We observed a considerable shift in the elasticities of both ecotypes when grown in the foreign environment. This suggests that the environment is a primary driver of the contributions to population growth, and aligns with differences in life history traits observed between genotypes, i.e., population growth is primarily driven by the early stages of the life cycle at low elevation, and by the late ones at high elevation. We interpret these results as indicative of cogradient plasticity of life history traits. This plasticity likely contributes to the fitness of the foreign ecotype at low and high elevation, but is not sufficient to match the fitness of the local ecotype. The evolution of plasticity is often observed in alpine plants and may be favored by strong daily and seasonal fluctuations in environmental conditions (Hassel *et al*., 2005; Ghalambor *et al*., 2007; Frei *et al*., 2014; Ensing & Eckert, 2019). It has been suggested that rapid shifts in selection imposed by climate change cannot be matched by an evolutionary response in perennial species (Anderson & Song, 2020; Anderson *et al*., 2019). However, adaptive plasticity may potentially mitigate the negative effects on population fitness and has been hypothesized to allow time for an evolutionary response to the novel selection pressure (Ghalambor *et al*., 2007; Fox *et al*., 2019; Vinton *et al*., 2022; West-Eberhard, 2003).

Our experiment highlights the utility of using demographic models to address questions related to adaptation of perennial plants. These approaches have only recently begun to be appreciated in adaptation studies (e.g., DeMarche *et al.,* 2016; Goebl *et al.,* 2022; but see Waser *et al*., 1985 for an older example) and represent an exciting opportunity to identify ecological drivers of ecotype formation (Wadgymar *et al*., 2022). By applying these approaches coupled with analyses of fitness trade-offs in populations growing in a reciprocal transplant experiment, we could reveal the contribution of life history traits to the evolution of elevational ecotypes in *D. carthusianorum*, and we could further assess how their expression responds to environmental variation. We found that plant development, life history traits and environmental heterogeneity contributed to patterns observed in fitness components at different stages of the life cycle. In particular, elevational adaptation appears to be driven by abiotic selection likely mediated by environmental stressors such as summer drought at low elevation, but also differences in resource availability linked to the length of the growing season at high elevation. We propose that future studies should aim to dissect the role of individual selective agents using field experiments, including manipulations, such as snow removal or drought treatments (Bushey *et al*., 2023). If used in combination with phenotypic selection analyses and forward genetic approaches such studies could establish a link between ecological knowledge on the effect of natural selection and the genetic basis of local adaptation.

## Acknowledgements

We are grateful to Köbi Graven, Stefan Hardegger and the Pfyn-Finges Natural Park for providing access to the sites used for the transplant gardens. We thank Maja Frei and Esther Zürcher for their support in the greenhouse work, Michael Gehrig for data collection from the sowing experiment, and members of the Plant Ecological Genetics group for help to set up the experiments. This project was supported by grants 31003A_160123 and 31003A_182675 from the Swiss National Science Foundation (SNSF) to AW.

## Author contributions

SF and AW designed the reciprocal transplant experiment, which SF and AP set up. AP and SF designed the establishment experiment, which AP set up. AP an UW collected the field data. AP performed all analyses. AP wrote the manuscript, which all authors revised.

## Data availability

