## Supporting information for "Life history traits mediate elevational adaptation in a perennial alpine plant"

**Supplementary figures**

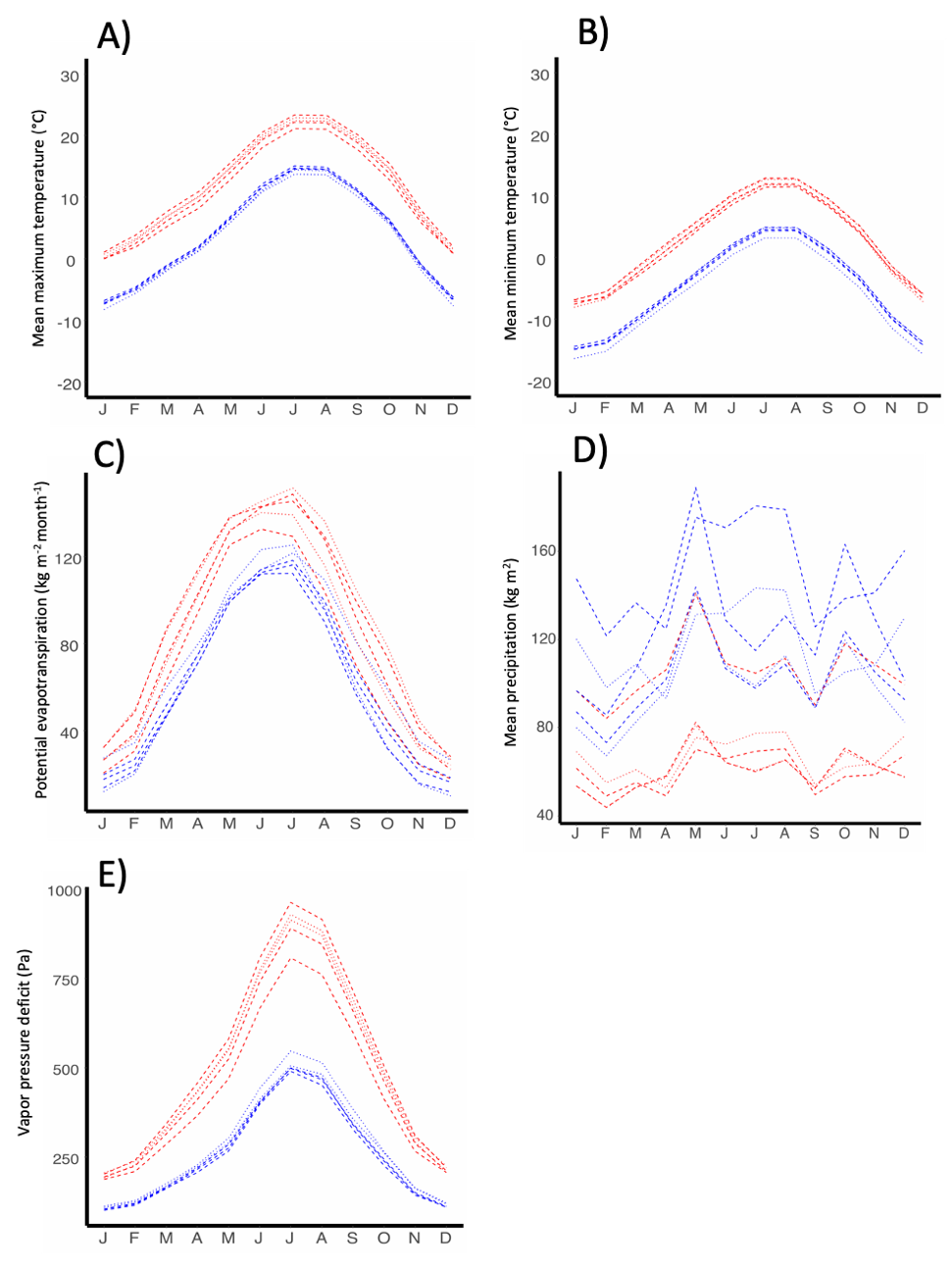

**Figure S1**. Mean monthly climatological estimates at low and high elevation transplant sites and at the original sites of the wild populations. A) mean maximum temperature, B) mean minimum temperature, C) Potential evapotranspiration (i.e., water loss through transpiration and evaporation), D) Precipitation, E) Vapor pressure deficit (i.e, difference between the amount of moisture in the air and the amount of moisture at saturation). Red and blue lines indicate low and high elevation, respectively. Dotted and dashed lines denote measures of the monthly means at our transplant sites and original sites of the wild populations, respectively. These estimates refer to the period 1980-2018, and were obtained from the Chelsa high-resolution database (Karger *et al.,* 2021).

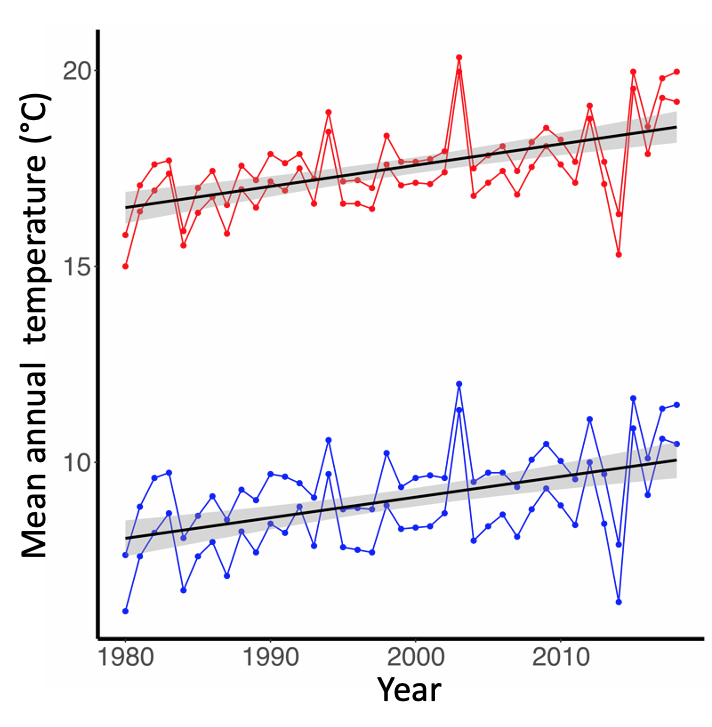

**Figure S2**. Change in mean annual temperature between 1980-2018 at low and high elevation transplant sites. Red and blue indicate low and high elevation, respectively. Estimates were calculated using the Chelsa high-resolution database (Karger *et al.,* 2021).

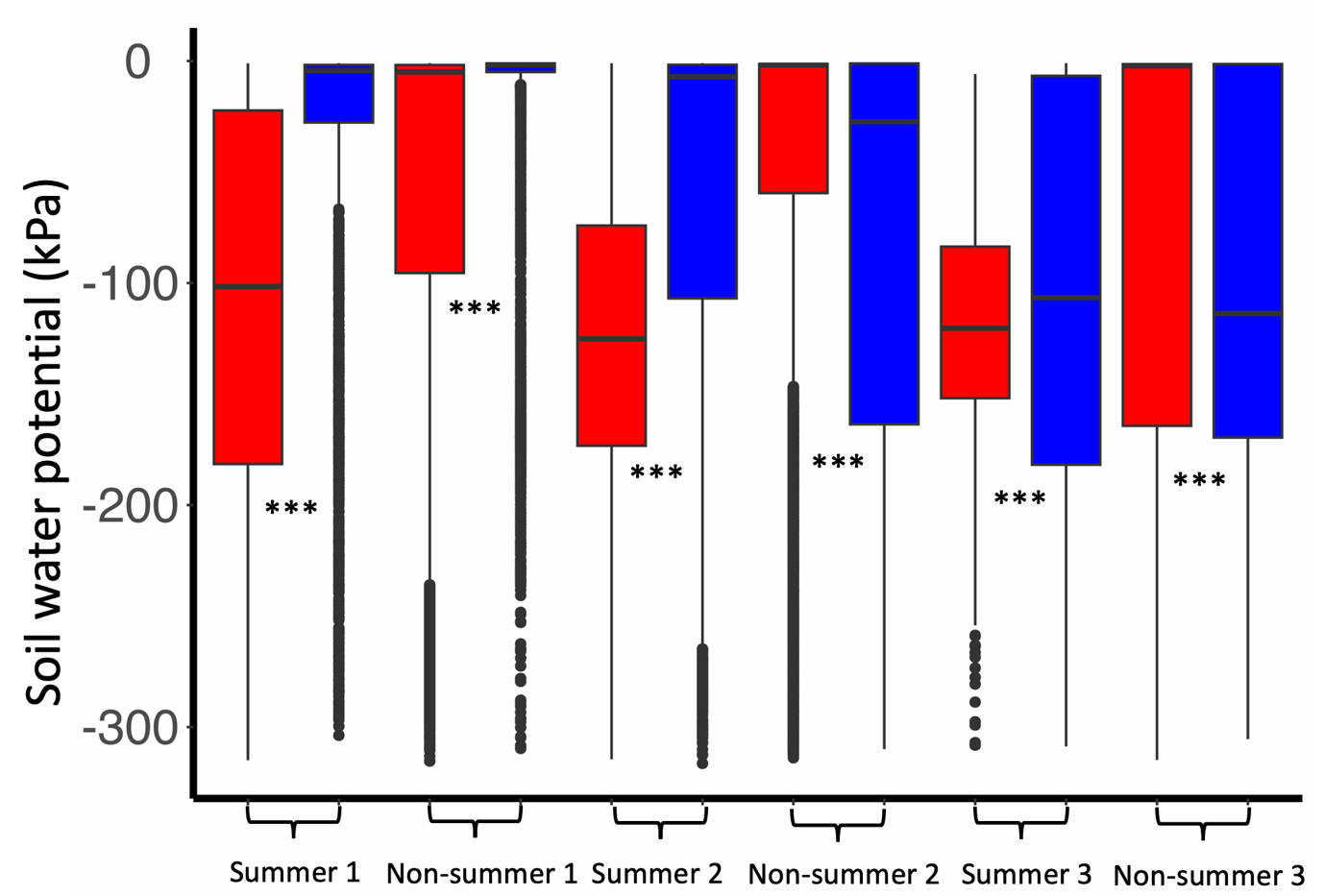

**Figure S3.** Soil water potential at low and high elevation transplant sites, A) over the duration of the experiment (2016-2018) and B) aggregated over the summers (i.e., months 6-8), rest of the year (i.e., months 1-5, 9-12) and year. Values are based on measurements from weather stations installed at each transplant site. Red and blue indicate low and high elevation, respectively. Significance of differences across elevation are reported (Wilcoxon rank-sum test; ***p<0.001).

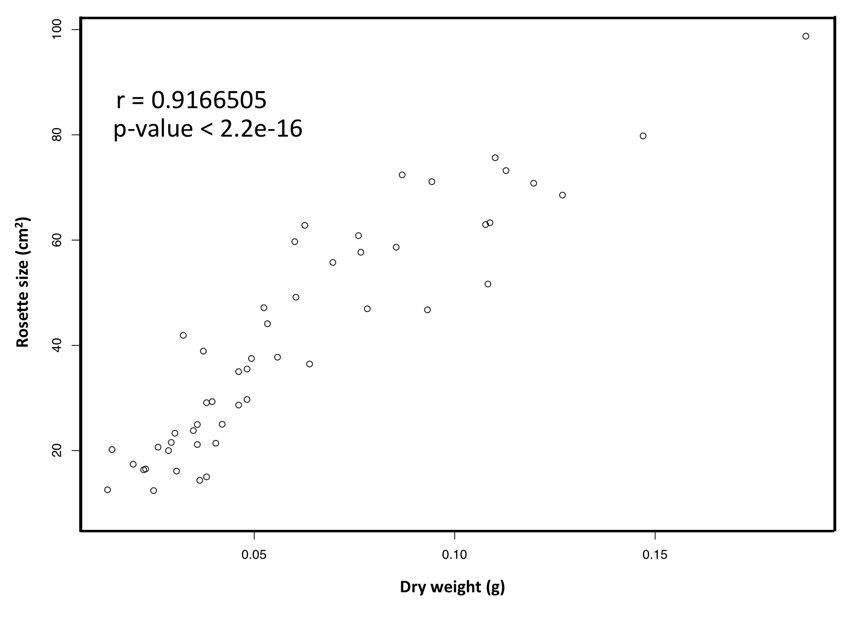

**Figure S4.** Correlation between rosette size (cm^2^) of and dry weight (g) of above ground tissue, based on data from 53 plants growing in a greenhouse facility (Lindau-Eschikon, Switzerland).

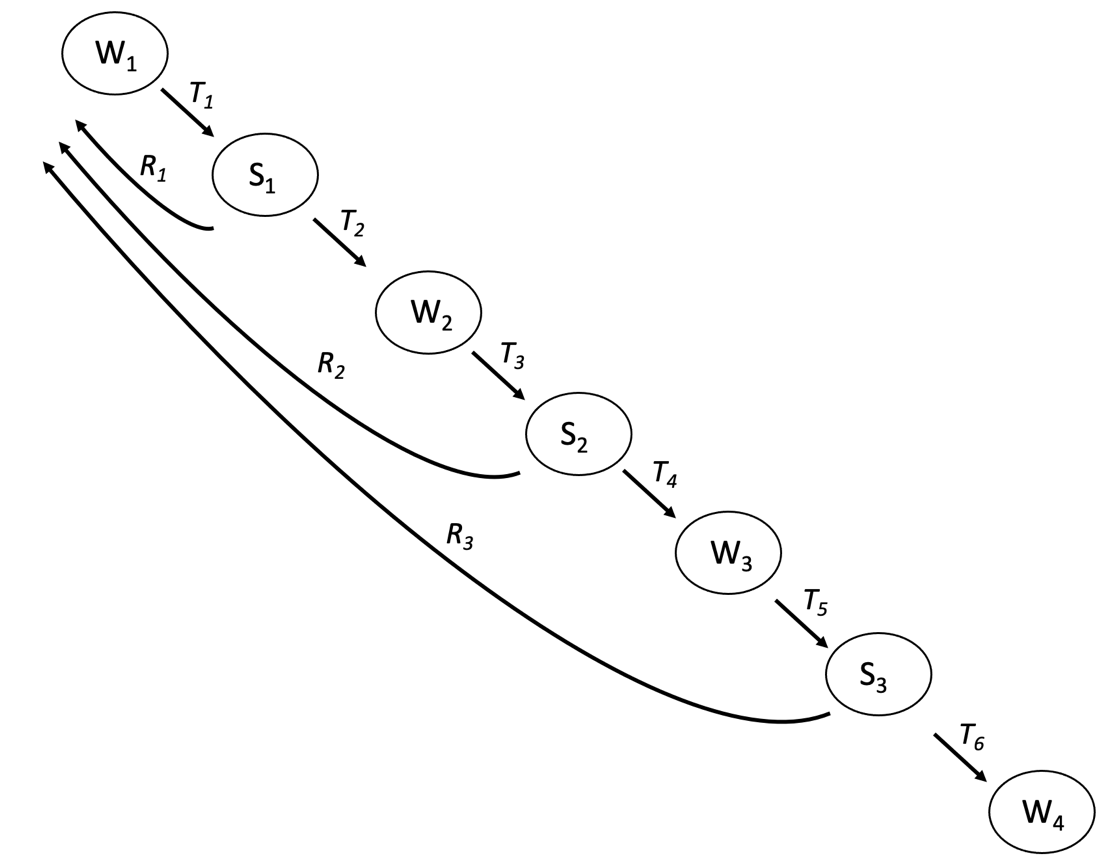

**Figure** **S5**. Age-classified life-cycle graph for *D. carthusianorum* used for the matrix population models. The life cycle is divided in subsequent summer (S_i_) and winter (W_i_) stages, represented by indexes in circles. The vital rates are inferred as transitions between the stages (i.e., survival T_i_ and reproduction R_i_).

**Supplementary tables**

**Table S1.** Transplant sites of the reciprocal transplant experiment with details on the sampled families used in the experiment.

| Transplant Site | Altitude | Meters above sea level | Coordinates: latitude, longitude | Number of maternal families | Mean nr. of seedlings per family ± SD |
| --- | --- | --- | --- | --- | --- |
| Leuk | Low | 890 | 46.27, 7.88 | 133 | 4.67 ± 3.48 |
| Zeneggen | Low | 930 | 46.31, 7.66 | 135 | 5.26 ± 3.63 |
| Findeln | High | 2120 | 46.01, 7.76 | 127 | 4.63 ± 2.88 |
| Oberu | High | 2150 | 46.35, 7.67 | 135 | 4.48 ± 2.98 |

**Table S2.** Low and high elevation populations used in the transplant experiment, with details on the sampled families.

| Population | Elevation | Meters above sea level | Coordinates: latitude, longitude | Number of maternal families | Mean nr. of seedlings per family ± SD |
| --- | --- | --- | --- | --- | --- |
| Unterstalden | Low | 754 | 46.26, 7.88 | 39 | 16.74 ± 12.93 |
| Niedergampel | Low | 688 | 46.31, 7.71 | 30 | 13.17 ± 14.77 |
| Grengiols | Low | 898 | 46.38, 8.11 | 23 | 16.48 ± 10.62 |
| Simplonpass | High | 2002 | 46.25, 8.03 | 20 | 18.10 ± 12.68 |
| Gibidumsee | High | 2211 | 46.26, 7.94 | 32 | 13.91 ± 10.58 |
| Faldumalp | High | 2029 | 46.38, 7.74 | 29 | 9.97 ± 7.05 |

**Table S3.** Outcome of the seed establishment experiment.

| **Model** | **Data** | **Mean establishment; SD** | **χ^2^; P-value** |
| --- | --- | --- | --- |
| Establishment ~ Genotype | Low Environment | Low: 0.17; 0.091  High: 0.14; 0.073 | 1.027; 0311 |
|  | High Environment | Low: 0.02; 0.028  High: 0.06; 0.06 | 2.778; 0.096 |

We modelled establishment as the response variable in generalized linear mixed effect models using a binomial error distribution. We tested the difference in establishment rates between genotypes in contrasting environments and obtained significance levels using likelihood ratio tests. We report mean establishment rates and their standard deviation (SD) per genotype extracted from the models, and χ^2^ p-values.

**Table S4**.Reproductive vital rates of elevational genotypes growing at low and high elevation transplant sites.

| **Test** | **S_1_: χ^2^; P-value** | **S_2_: χ^2^; P-value** | **S_3_: χ^2^; P-value** |
| --- | --- | --- | --- |
| **Flowering probability** |  |  |  |
| *Genotype X Environment* | **5.116;** **0.024** | **39.008; <0.001** | **46.8; <0.001** |
| *Local vs. foreign* | **S_1_: Estimate; SE; P-value** | **S_2_: Estimate; SE; P-value** | **S_3_: Estimate; SE; P-value** |
| Low Environment | 0.573; 0.430; 0.458 | **0.218;** **0.063; <.0001** | **0.325;** **0.106; <0.001** |
| High Environment | 0.965; 0.721; 0.962 | 1.320; 0.353; 0.300 | **3.218; 1.132; <0.034** |
| *Home vs. away* |  |  |  |
| Low genotype | **0.408; 0.137; 0.008** | **0.222;** **0.115; 0.004** | **0.012; 0.012; <0.001** |
| High genotype | 0.688; 0.222; 0.246 | 1.347; 0.708; 0.571 | **0.120; 0.120; 0.034** |
| **Seed output** | **S_1_: χ^2^; P-value** | **S_2_: χ^2^; P-value** | **S_3_: χ^2^; P-value** |
| *Genotype X Environment* | **87.829; <0.001** | **69.125; <0.001** | **78.41; <0.001** |
| *Local vs. foreign* | **S_1_: Estimate; SE; P-value** | **S_2_: Estimate; SE; P-value** | **S_3_: Estimate; SE; P-value** |
| Low Environment | 0.835; 0.233; 0.518 | **0.481;** **0.165;** **0.033** | -0.70;2 0.509; 0.168 |
| High Environment | 0.609; 0.171; 0.077 | 1.407; 0.482; 0.319 | -0.99; 0.510; 0.052 |
| *Home vs. away* |  |  |  |
| Low genotype | **0.689 ; 0.126; 0.042** | 0.628; 0.269; 0.277 | -3.24; 2.179; 0.137 |
| High genotype | **0.502; 0.092; <.0001** | 0.839; 0.790; 0.156 | -3.53; 2.179; 0.105 |

We modelled flowering probability and seed output by implementing them as response variables in generalized linear mixed effect models. Flowering probability was modelled with a binomial error distribution and seed output using a zero-inflated Poisson error distributions. We tested for genotype by environment interactions and differential performance of the genotypes according to the local vs. foreign and home vs. away criteria of local adaptation. We tested the significance of the interaction between elevation and genotype using likelihood ratio tests and report χ^2^ and p-values. Local vs. foreign and home vs. away contrasts were estimated using pairwise contrasts. S_i_ indicates the summers and significant results are in bold.

**Table S5.** Size of elevational genotypes growing at low and high elevation transplant sites.

| **Test** | **χ^2^; P-value** | | | | | |
| --- | --- | --- | --- | --- | --- | --- |
| **Size** | W_1_ | S_1_ | W_2_ | S_2_ | S_3_ | W_4_ |
| *Genotype X  Environment* | 1.888; 0.171 | **10.226; 0.001** | **5.285; 0.022** | **47.695; <0.001** | **44.354; <0.001** | 0.807; 0.369 |
| *Local vs. foreign* |  |  | **Estimate; SE; P-value** | |  |  |
| Low Environment | -1.079; 0.427; 0.064 | -0.841; 0.360; 0.077 | **-1.305; 0.363; 0.019** | **-0.947; 0.197; 0.005** | **-0.62;** **0.165;** **0.013** | **-0.326; 0.137; 0.032** |
| High Environment | **-1.197;** **0.427; 0.048** | -0.559; 0.358; 0.192 | **-1.022; 0.357; 0.044** | -0.270; 0.192; 0.226 | -0.019; 0.160; 0.8909 | **-0.436; 0.104;** **0.009** |
| *Home vs. away* |  |  |  |  |  |  |
| Low genotype | -0.103; 0.238; 0.704 | **0.796;** **0.199; 0.046** | 0.686; 0.318; 0.154 | -0.258; 0.573; 0.695 | -1.367; 0.597; 0.148 | -0.207; 0.278; 0.528 |
| High genotype | -0.221; 0.237; 0.444 | **1.079;** **0.196; 0.027** | 0.970; 0.318; 0.084 | 0.418; 0.573; 0.540 | -0.762; 0.597; 0.329 | -0.317; 0.281; 0.364 |

We modelled size by implementing it as response variables linear mixed effect models with a Gaussian error distribution. We tested for genotype by environment interactions and differential performance of the genotypes according to the local vs. foreign and home vs. away criteria of local adaptation. We tested the significance of the interaction between elevation and genotype using likelihood ratio tests and report χ^2^ and p-values. Local vs. foreign and home vs. away contrasts were estimated using pairwise contrasts. W_i_ and S_i_ indicate winter and summer, respectively. Significant results are in bold.

**Table S6**. Cumulative survival of elevational genotypes growing at low and high elevation transplant sites.

| **Test** | **Coefficients** | **Hazard ratio** | **P-values** |
| --- | --- | --- | --- |
| *Genotype X Environment* | **-1.732** | **0.177** | **<0.001** |
| *Local vs. foreign* |  |  |  |
| Low Environment | **-1.005** | **0.349** | **0.022** |
| High E Environment | **0.762** | **2.143** | **0.003** |
| *Home vs. away* |  |  |  |
| Low genotype | 0.017 | 1.017 | 0.955 |
| High genotype | **1.805** | **6.081** | **<0.001** |

We modelled survival throughout the experiment using Cox proportional hazard models and tested for genotype by environment interactions and differential performance of the genotypes according to the local vs. foreign and home vs. away criteria of local adaptation. Significant results are in bold.

**Table S7.** Single time point survival of elevational genotypes growing at low and high elevation transplant sites.

| **Test** | **χ^2^; P-value** | | | | | |  |
| --- | --- | --- | --- | --- | --- | --- | --- |
| **Survival** | W_1_ | S_1_ | W_2_ | S_2_ | W_3_ | S_3_ | W_4_ |
| *Genotype X Environment* | **33.573; <0.001** | **13.471; <0.001** | **22.744; <0.001** | 1.470; 0.225 | **3.606; 0.058** | 0.046; 0.838 | 3.342; 0.068 |
| *Local vs. foreign* | |  |  | **Estimate; SE; P-value** |  |  |  |
| Low Environment | 0.549; 0.251; 0.189 | **0.166;** **0.082; <0.001** | 0.794; 0.213; 0.390 | **0.194; 0.082; <0.001** | 0.235; 0.217; 0.118 | 0.853; 0.236; 0.565 | **0.422;** **0.178; 0.041** |
| High Environment | **10.755; 5.29 <0.001** | 0.674; 0.371; 0.474 | **5.208;** **1.814; <0.001** | 0.380; 0.203; 0.070 | 1.644; 1.041; 0.432 | 0.944; 0.389; 0.889 | 1.153; 0.405; 0.685 |
| *Home vs. away* | |  |  |  |  |  |  |
| Low genotype | **0.140; 0.091; 0.002** | **2.706; 1.111; 0.015** | 1.009; 0.356; 0.980 | 3.383; 2.833; 0.146 | 0.317; 0.369; 0.324 | 3.635; 2.608; 0.072 | 1.009; 1.255; 0.994 |
| High genotype | 2.749; 1.884; 0.140 | **10.978;** **3.877; <0.001** | **6.614;** **2.775; <0.001** | **6.638; 4.995;** **0.012** | 2.220; 2.334; 0.448 | **4.022; 2.814; 0.047** | 2.757; 3.387; 0.409 |

We modelled survival at specific stages of the life cycle by implementing it as response variable generalized linear mixed effect models with a binomial error distribution. We tested for genotype by environment interactions and differential performance of the genotypes according to the local vs. foreign and home vs. away criteria of local adaptation. We tested the significance of the interaction between elevation and genotype using likelihood ratio tests and report χ^2^ and p-values. Local vs. foreign and home vs. away contrasts were estimated using pairwise contrasts. W_i_ and S_i_ indicate winter and summer, respectively. Significant results are in bold.

**Table S8**. Population growth rate and LTRE at the two elevational transplant environments.

| **Environment** | **Lambda, high genotype** | **Lambda, low genotype** | **T_1_** | **R_1_** | **T_2_** | **T_3_** | **R_2_** | **T_4_** | **T_5_** | **R_3_** | **T_6_** |
| --- | --- | --- | --- | --- | --- | --- | --- | --- | --- | --- | --- |
| High | 0.72 (0.67, 0.77) | 0.46 (0.41, 0.53) | -0.019 | -0.044 | -0.001 | -0.010 | -0.032 | 0.005 | <-0.001 | -0.116 | 0 |
| Low | 1.50 (1.35, 1.70) | 2.03 (1.77, 2.40) | -0.045 | -0.408 | -0.002 | -0.012 | -0.025 | 0.003 | <-0.001 | -0.031 | 0 |

Population growth rate of the elevational genotypes growing in the low and high elevation transplant environments, mean values based on 20 000 bootstrap replicates and 95% bias corrected confidence intervals are reported. Results of the LTRE show the contribution of the vital rates of the foreign genotype to population growth rate in the two transplant environments. Survival and reproductive vital rates throughout the life cycle are indicated by T_i_ and R_i_, respectively. Mean estimates based on 20 000 bootstrap replicates are reported.

**Table S9.** Trade-offs between plant size and flowering probability.

| **Model** | | **S_1_: χ^2^; P-value** | **S_2_: χ^2^; P-value** | **S_3_: χ^2^; P-value** |  |  |  |
| --- | --- | --- | --- | --- | --- | --- | --- |
| Flowering ~ Plant size X Genotype X Environment | | **28.054; <0.001** | **5.462; 0.019** | 0.181; 0.671 |  |  |  |
| Flowering ~ Plant size X Genotype | | **Low E: 50.106; <0.001, High E: 19.529; <0.001** |  |  |  |  |  |
| **Time point** | **Environment** | **Genotype** | **Trend, plant size** | **SE** | **df** | **z.ratio** | **p-value** |
| S_1_ | **Low** | **Low** | **1.08E-03** | **3.88E-04** | **Inf** | **2.797** | **0.005** |
|  | **Low** | **High** | **8.81E-03** | **1.19E-03** | **Inf** | **7.426** | **<.001** |
|  | **High** | **Low** | **1.14E-03** | **2.10E-04** | **Inf** | **5.562** | **<.001** |
|  | **High** | **High** | **2.63E-03** | **2.73E-04** | **Inf** | **9.615** | **<.001** |
| S_2_ | **Low** | **Low** | 1.31E-04 | 1.42E-04 | Inf | 1.597 | 0.110 |
|  | **Low** | **High** | **1.18E-03** | **1.57E-04** | **Inf** | **3.162** | **0.002** |
|  | **High** | **Low** | **6.73E-04** | **1.55E-04** | **Inf** | **4.337** | **<.0001** |
|  | **High** | **High** | **9.46E-04** | **1.42E-04** | **Inf** | **6.683** | **<.0001** |
| S_3_ | **Low** | **Low** | **3.90E-04** | **1.25E-04** | **Inf** | **3.121** | **0.002** |
|  | **Low** | **High** | **1.12E-03** | **2.47E-04** | **Inf** | **4.518** | **<.0001** |
|  | High | Low | **1.20E-03** | **3.81E-04** | **Inf** | **3.148** | **0.002** |
|  | **High** | **High** | **2.16E-03** | **2.88E-04** | **Inf** | **7.518** | **<.0001** |

We tested for trade-offs between plant size and flowering throughout the experiment by modelling flowering probability as a function of plant size using generalized linear mixed effect models. We tested for three way interactions between genotypes, plant size and the transplant environment and two-way interactions between genotype and plant size within the low and high elevation transplant environments, respectively as well as trends within experimental genotypes. We tested the significance of the interactions using likelihood ratio tests and report χ^2^ and p-values. S_i_ denotes the summer. Significant results are in bold.

**Table S10.** Trade-offs between plant size and survival probability.

| **Model** | | **W_1_: χ^2^; P-value** | **S_1_: χ^2^; P-value** | **W_2_: χ^2^; P-value** | **S_2_: χ^2^; P-value** | **S_3_: χ^2^; P-value** | **W_4_: χ^2^; P-value** |
| --- | --- | --- | --- | --- | --- | --- | --- |
| Survival~ Plant size X Genotype X Environment | | 0.379; 0.538 | **10.693; 0.001** | 2.049; 0.152 | **5.818; 0.016** | 0.298; 0.585 | 0.654; 0.419 |
| Survival ~ Plant size X Genotype | |  | **Low E: 4.736; 0.03**, **High E: 6.203; 0.013** | **Low E: 5.314; 0.021** , High E: 2.130; 0.144 |  |  |  |
| **Time point** | **Environment** | **Genotype** | **Trend, plant size** | **SE** | **df** | **z.ratio** | **p-value** |
| Survival W_1_ | **Low** | **Low** | 5.31E-04 | 4.82E-04 | Inf | 1.103 | 0.27 |
|  | **Low** | **High** | **3.51E-03** | **1.07E-03** | **Inf** | **3.28** | **0.001** |
|  | **High** | **Low** | **1.71E-3** | **3.79E-04** | **Inf** | **4.504** | **<0.0001** |
|  | High | High | 6.96E-03 | 3.76E-03 | Inf | 1.851 | 0.0642 |
| Survival S_1_ | Low | Low | 8.44E-04 | 5.36E-04 | **Inf** | 1.576 | 0.115 |
|  | **Low** | **High** | **2.29E-03** | **7.81E-04** | **Inf** | **3.735** | **0.0002** |
|  | **High** | **Low** | **6.71E-03** | **1.76E-03** | **Inf** | **3.805** | **0.0001** |
|  | **High** | **High** | **2.53E-03** | **7.24E-03** | **Inf** | **3.496** | **0.0005** |
| Survival W_2_ | Low | Low | 3.18E-04 | 4.20E-04 | **Inf** | 0.758 | 0.449 |
|  | **Low** | **High** | **-2.04E-3** | **9.25E-04** | **Inf** | **-2.06** | **0.027** |
|  | **High** | **Low** | **6.41E-04** | **2.87E-04** | **Inf** | **2.235** | **0.025** |
|  | High | High | -1.08E-04 | 3.93E-04 | Inf | -0.274 | 0.784 |

We tested for trade-offs between plant size and survival throughout the experiment by modelling survival probability as a function of plant size using generalized linear mixed effect models. We tested for three way interactions between genotypes, size and the transplant environment and two-way interactions between genotype and size within the low and high elevation transplant environments, respectively as well as trends within experimental genotypes. We tested the significance of the interactions using likelihood ratio tests and report χ^2^ and p-values. W_i_ and S_i_ indicate winter and summer, respectively. Significant results are in bold.

| **Time point** | **Environment** | **Genotype** | **Trend, plant size** | **SE** | **df** | **z.ratio** | **p-value** |
| --- | --- | --- | --- | --- | --- | --- | --- |
| Survival S_2_ | Low | Low | 2.95E-04 | 2.98E-04 | Inf | 0.990 | 0.322 |
|  | Low | High | 1.20E-03 | 8.46E-04 | Inf | 1.502 | 0.133 |
|  | **High** | **Low** | **1.68E-02** | **7.08E-03** | **Inf** | **2.377** | **0.018** |
|  | **High** | **High** | **4.81E-03** | **1.35E-03** | **Inf** | **3.554** | **0.0004** |
| Survival S_3_ | Low | Low | 2.35E-04 | 1.41E-04 | Inf | 1.665 | 0.096 |
|  | Low | High | 3.34E-04 | 2.82E-04 | Inf | 1.182 | 0.237 |
|  | High | Low | 1.67E-03 | 1.21E-03 | Inf | 1.383 | 0.167 |
|  | High | High | 1.0E-03 | 7.06E-04 | Inf | 1.427 | 0.154 |
| Survival W_4_ | Low | Low | 1.33E-04 | 1.88E-04 | Inf | 0.707 | 0.480 |
|  | Low | High | 1.78E-04 | 3.54E-04 | Inf | 0.503 | 0.615 |
|  | **High** | **Low** | **4.05E-03** | **1.30E-03** | **Inf** | **3.119** | **0.002** |
|  | **High** | **High** | **2.74E-03** | **1.00E-03** | **Inf** | **2.726** | **0.006** |

**Table S10.** Continued.

**Table S11.** Trade-offs between survival and flowering probability.

| **Model** | | | **S_1_: χ^2^; P-value** | | **W_2_: χ^2^; P-value** | | **S_2_: χ^2^; P-value** | **W_3_: χ^2^; P-value** | | **S_3_: χ^2^; P-value** | | **W_4_: χ^2^; P-value** | |
| --- | --- | --- | --- | --- | --- | --- | --- | --- | --- | --- | --- | --- | --- |
| Survival ~ Flowering X Genotype X Environment | | | **8.42; 0.004** | | **8.81; 0.003** | | 0.08; 0.77 | 0.0; 0.99 | | 0.01;0.94 | | 0.98; 0.32 | |
| Survival ~ Flowering X Genotype | | | **Low E:** **5.18; 0.02** , **High E: 4.21; 0.04** | | Low E: 8.14; 0.004**, High E: 2.42; 0.12** | |  |  | |  | |  | |
| **Time point** | **Environment** | **Genotype** | | **Contrast, flowering** | | **Estimate** | | **SE** | **df** | | **z.ratio** | | **p-value** |
| Survival S_1_ | **Low** | **Low** | | **yes - no** | | **3.272** | | **0.431** | **Inf** | | **7.596** | | **<0.001** |
|  | **Low** | **High** | | **yes - no** | | **1.619** | | **0.237** | **Inf** | | **6.835** | | **<0.001** |
|  | **High** | **Low** | | **yes - no** | | **3.447** | | **0.531** | **Inf** | | **6.487** | | **<0.001** |
|  | **High** | **High** | | **yes - no** | | **3.321** | | **0.729** | **Inf** | | **4.556** | | **<0.001** |
| Survival W_2_ | Low | Low | | yes - no | | -0.833 | | 0.51 | Inf | | -1.632 | | 0.6 |
|  | **Low** | **High** | | **yes - no** | | **-2.571** | | **0.728** | **Inf** | | **-3.533** | | **0.005** |
|  | High | Low | | yes - no | | -0.645 | | 0.359 | Inf | | -1.796 | | 0.479 |
|  | High | High | | yes - no | | 0.894 | | 0.576 | Inf | | 1.553 | | 0.657 |

We tested for trade-offs between survival and flowering throughout the experiment by modelling survival probability as a function of flowering using generalized linear mixed effect models. We tested for three way interactions between genotypes, flowering and the transplant environment and two-way interactions between genotype and survival within the low and high elevation transplant environments, respectively as well as trends within experimental genotypes. We tested the significance of the interactions using likelihood ratio tests and report χ^2^ and p-values. W_i_ and S_i_ indicate winter and summer, respectively. Significant results are in bold.

**Table S11**. Continued.

| **Time point** | **Environment** | **Genotype** | **Contrast, flowering** | **Estimate** | **SE** | **df** | **z.ratio** | **p-value** |
| --- | --- | --- | --- | --- | --- | --- | --- | --- |
| Survival W_2_ | Low | yes | low-high | 0.509 | 0.27 | Inf | 1.882 | 0.419 |
|  | Low | no | low-high | -1.23 | 0.849 | Inf | -1.448 | 0.73 |
|  | **High** | **yes** | **low-high** | **-2.413** | **0.497** | **Inf** | **-4.86** | **<0.001** |
|  | High | no | low-high | -0.874 | 0.469 | Inf | -1.865 | 0.43 |
| Survival S_2_ | Low | Low | yes - no | 18.593 | 3563.284 | Inf | 0.005 | 1 |
|  | Low | High | yes - no | 19.56 | 6070.393 | Inf | 0.003 | 1 |
|  | High | Low | yes - no | 17.494 | 6098.544 | Inf | 0.003 | 1 |
|  | High | High | yes - no | 18.13 | 4738.979 | Inf | 0.004 | 1 |
| Survival W_3_ | Low | Low | yes - no | -0.879 | 1.243 | Inf | -0.707 | 0.991 |
|  | Low | High | yes - no | -0.16 | 1.173 | Inf | -0.136 | 0.999 |
|  | High | Low | yes - no | 0.03 | 1.117 | Inf | 0.027 | 1 |
|  | High | High | yes - no | 17.024 | 63.634 | Inf | 0.268 | 0.999 |
| Survival S_3_ | Low | Low | yes - no | 0.608 | 0.384 | Inf | 1.582 | 0.638 |
|  | Low | High | yes - no | 0.454 | 0.38 | Inf | 1.196 | 0.877 |
|  | High | Low | yes - no | -0.487 | 1.046 | Inf | -0.465 | 0.999 |
|  | High | High | yes - no | 0.465 | 0.751 | Inf | 0.62 | 0.997 |
| Survival W_4_ | Low | Low | yes - no | -0.874 | 0.65 | Inf | -1.344 | 0.797 |
|  | Low | High | yes - no | 0.226 | 0.53 | Inf | 0.428 | 0.999 |
|  | High | Low | yes - no | -1.19 | 1.04 | Inf | -1.145 | 0.899 |
|  | High | High | yes - no | 0.116 | 0.988 | Inf | 0.117 | 0.999 |

**Table S12.** Influence of specific vital rates on population growth rate at the two transplant environments.

| **Environment** | **Genotype** | **T_1_** | **R_1_** | **T_2_** | **T_3_** | **R_2_** | **T_4_** | **T_5_** | **R_3_** | **T_6_** |
| --- | --- | --- | --- | --- | --- | --- | --- | --- | --- | --- |
| High | High | 0.254 (0.226, 0.287) | 0.112 (0.074, 0.162) | 0.141 (0.124, 0.156) | 0.141 (0.124, 0.156) | 0.036 (0.02, 0.064) | 0.105 (0.082, 0.125) | 0.105 (0.082, 0.125) | 0.105 (0.082, 0.125) | 0 |
| High | Low | 0.243 (0.208, 0.29) | 0.111 (0.06, 0.182) | 0.132 (0.109, 0.148) | 0.132 (0.109, 0.148) | 0.007 (0.002, 0.019) | 0.125 (0.101, 0.144) | 0.125 (0.101, 0.144) | 0.125 (0.101, 0.144) | 0 |
| Low | High | 0.432 (0.381, 0.469) | 0.398 (0.321, 0.454) | 0.034 (0.016, 0.06) | 0.034 (0.016, 0.06) | 0.0005 (0, 0.002) | 0.034 (0.015, 0.059) | 0.034 (0.015, 0.059) | 0.034 (0.015, 0.059) | 0 |
| Low | Low | 0.454 (0.416, 0.483) | 0.426 (0.366, 0.471) | 0.028 (0.011, 0.05) | 0.028 (0.011, 0.05) | 0.01 (0.005, 0.019) | 0.018 (0.006, 0.035) | 0.018 (0.006, 0.035) | 0.018 (0.006, 0.035) | 0 |

Influence of specific vital rates on population growth rates extracted from the matrix population models of the elevational genotypes growing in the low and high elevation transplant environments expressed as elasticities. Survival and reproductive vital rates throughout the life cycle are indicated by T_i_ and R_i_, respectively, Mean values are based on 20 000 bootstrap replicates and 95% bias corrected confidence intervals are reported.

**Table S13.** Stable age distribution of the elevational genotypes growing in the low and high elevation transplant environments.

| **Environment** | **Genotype** | **W_1_** | **S_1_** | **W_2_** | **S_2_** | **W_3_** | **S_3_** | **W_4_** |
| --- | --- | --- | --- | --- | --- | --- | --- | --- |
| High | High | 0.085 (0.065, 0.109) | 0.116 (0.093, 0.142) | 0.099 (0.085, 0.115) | 0.133 (0.121, 0.147) | 0.145 (0.133, 0.159) | 0.197 (0.173, 0.223) | 0.224 (0.182, 0.27) |
| High | Low | 0.015 (0.007, 0.027) | 0.027 (0.016, 0.045) | 0.035 (0.023, 0.053) | 0.066 (0.049, 0.088) | 0.118 (0.099, 0.142) | 0.251 (0.228, 0.279) | 0.488 (0.405, 0.559) |
| Low | High | 0.501 (0.448, 0.56) | 0.303 (0.283, 0.327) | 0.104 (0.085, 0.122) | 0.048 (0.035, 0.062) | 0.023 (0.014, 0.032) | 0.015 (0.008, 0.023) | 0.006 (0.003, 0.01) |
| Low | Low | 0.594 (0.54, 0.659) | 0.283 (0.253, 0.311) | 0.074 (0.056, 0.092) | 0.032 (0.02, 0.044) | 0.01 (0.005, 0.016) | 0.005 (0.002, 0.009) | 0.001 (0.001, 0.003) |

Winter and summer stages are represented by W_i_ and S_i_, respectively. Values are based on 20 000 bootstrap replicates and 95% bias corrected confidence intervals are reported.
